# Overriding bioprocess perturbations with a cell–machine interface for reliable microbial stress-response control

**DOI:** 10.1101/2025.11.21.689713

**Authors:** Mathéo Delvenne, Juan Andres Martinez, Cees Haringa, Henk Noorman, Steven Minden, Ralf Takors, Frank Delvigne

## Abstract

Controlling cell population dynamics and phenotypic diversification is a key objective in systems and synthetic biology, particularly for ensuring uniform responses from engineered gene circuits. While cell-machine interfaces have been employed to modulate host–gene circuit interactions, environmental perturbations typical of industrial bioreactor conditions remain underexplored. In this study, we investigate the impact of such perturbations on the general stress response in *Escherichia coli* and *Saccharomyces cerevisiae*. Using scale-down bioreactor experiments, we evaluate the performance of the Segregostat, a real-time control system that leverages automated flow cytometry to induce dynamic nutrient shifts. The Segregostat achieves robust stress response control, even under severe perturbations such as extended residence times in a two-compartment reactor. We hypothesize that this robustness arises from the system’s ability to amplify host-compatible fluctuations beyond bioreactor-induced perturbations. Our findings highlight the importance of integrating environmental factors into control strategies for reliable gene circuit behavior in industrial bioprocessing environments.

## Introduction

Systems and synthetic biology have significantly advanced our ability to design and control synthetic or natural gene circuits, yet most work has focused on their interactions with the cellular host ^1,2^, addressing challenges such as metabolic burden ^3^, switching dynamics ^4^, and population heterogeneity ^5^. Meanwhile, bioprocess engineering has concentrated on optimizing the interplay between cells and their physical environment within bioreactors ^6,7^. On top of these parallel advances, progress in fields such as industrial biotechnology, therapeutic protein production, and biosensing increasingly depends on bridging these two domains to create scalable and controllable bioprocesses ^8^. A key step toward this integration is the development of robust frameworks that account for the full spectrum of interactions i.e., between gene circuits, host physiology, and bioreactor conditions. Recent innovations have introduced cell–machine interfaces capable of connecting microbial populations to external computers and controllers, enabling real-time modulation of gene expression ^9–11^. However, these technologies remain largely untested under the dynamic and often harsh conditions relevant to industrial-scale bioprocessing ^12^.

In this work, we address this gap by evaluating the Segregostat, a real-time control platform for microbial populations ^13^, under scale-down bioreactor conditions. This approach enables us to assess the robustness, responsiveness, and scalability of cell–machine interfaces, paving the way for their deployment in industrial biotechnology applications. More specifically, we employed the Segregostat to modulate the general stress response in *S. cerevisiae* and *E. coli*. These regulatory networks, previously shown to generate substantial population heterogeneity ^14–16^, are of particular interest for bioprocess optimization ^17,18^, where controlling their activity could mitigate growth arrest and reduce the metabolic burden associated with their activation ^19,20^. Under standard laboratory-scale bioreactor conditions, the Segregostat consistently reduced phenotypic heterogeneity, reflecting cell-to-cell variability in stress response, in both *E. coli* and *S. cerevisiae*. The magnitude of this reduction scaled proportionally with the metabolic burden imposed by activation of the target gene circuit, in agreement with previous observations ^5^. We next implemented a two-compartment scale-down reactor (SDR) to reproduce the concentration gradients characteristic of poorly mixed large-scale bioreactors ^8,21^. Even under these highly heterogeneous conditions, the Segregostat maintained robust population-level control, demonstrating resilience to severe mixing limitations.

## Results

### Segregostat reduces cell-to-cell heterogeneity (entropy) in stress response at laboratory scale

A variety of cell–machine interfaces are now available to enable direct communication with targeted cell populations and to monitor and/or control their physiological states ^9–11,13,22–25^. Among them, we utilized the Segregostat, a platform that combines automated flow cytometry (AFC) with a feedback control loop, to regulate the general stress response in bacterial and yeast populations (**Figure 1 A**). Under chemostat conditions, nutrient limitation triggers the transition to stress states. This transition can be counteracted by timely glucose pulses. In our setup, glucose was automatically pulsed into the bioreactor whenever single-cell fluorescence measurements indicated that more than 50% of the population exceeded a defined stress threshold (**Figure 1 A-B**). This approach created alternating phases of feast (glucose pulsing, promoting growth) and famine (no pulsing, promoting nutrient limitation and stress responses), ultimately driving coordinated and synchronized gene expression within the population ^5^.

**Figure 1:**
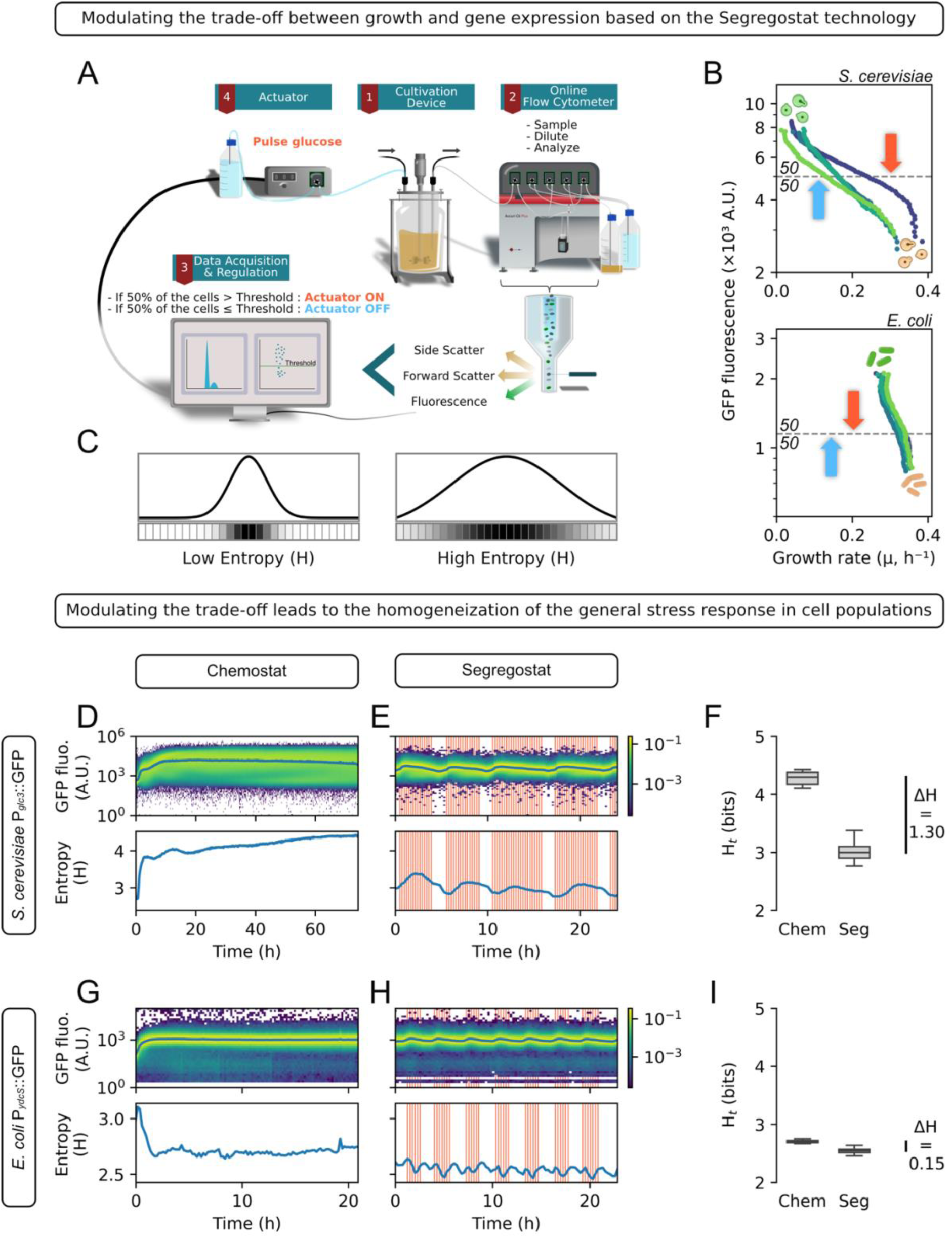
Synchronization and homogenization of stress-related genes based on the Segregostat technology. **A** Schematic representation of the Segregostat setup. **B** Control loop of the Segregostat and illustration of the trade-off between growth and stress-related gene expression in *E. coli* and *S. cerevisiae*. The horizontal gray dashed line indicates the fluorescence threshold. Red arrow indicates activation of glucose pulse, while blue once indicates its inactivation. The colored dots indicate different replicates for computing the trade-off, adapted from Delvenne *et al.* (2025). **C** Schematic explanation of population heterogeneity quantification by Shannon entropy (H). This measure comes from information theory and is computed from the distribution of the population across discrete bins. **D-E** Fluorescence time-scatter plots and corresponding entropy dynamics of *S. cerevisiae* P*_glc3_*::GFP in continuous chemostat cultivation at D = 0.1 h⁻¹ (**D**) and under Segregostat control (Threshold = 3000 A.U., 0.4 g glucose/pulse) (**E**). Glucose pulses are marked by red vertical lines. **G-H** Fluorescence time-scatter plots and corresponding entropy dynamics of *E. coli* P*_ydcS_*::GFP in continuous chemostat cultivation at D = 0.3 h⁻¹(**G**) and under Segregostat control (Threshold = 1150 A.U., 1 g glucose/pulse) (**H**). Glucose pulses are marked by red vertical lines. **F–I** Basal entropy of *S. cerevisiae* (**F**) and *E. coli* (**I**), calculated from the fluorescence distribution of the population over time, represented as boxplots. For chemostat conditions, entropy is computed from 3 residence times onward until the end of cultivation. For Segregostat conditions, only the last 3–4 oscillations are considered.

Previously, we demonstrated that the degree of cell-to-cell heterogeneity in stress-related gene expression, quantified here using Shannon entropy (**Figure 1 C**), is primarily driven by the fitness or switching cost, defined as the reduction in growth rate experienced by cells that activate stress responses ^5^. By exploiting the alternation of stress induction and relaxation phases generated by the Segregostat, we were able to quantify the trade-off between growth and the expression of a stress-related gene (**Figure 1 B**) ^26^. Interestingly, this fitness cost differs significantly between *Escherichia coli* and *Saccharomyces cerevisiae* (**Figure 1 B**) ^26^. In *S. cerevisiae*, the fitness cost is considerable and leads to a highly heterogeneous activation of the general stress response under chemostat conditions, as reflected by elevated entropy levels (**Figure 1 D**). Functionally, this elevated entropy enables the population to maintain an average growth rate above the dilution rate, even though individual cells may incur significant growth penalties. This population-level resolution of the trade-off between growth and gene expression has been previously described as the Fitness–Entropy Compensation effect ^26^. This effect is particularly pronounced when monitoring the *P_glc3_::GFP* transcriptional reporter at low dilution rates, where stress responses are robustly induced ^5,26^ (**Figure 1 D**, **Supp. Note 1**). By contrast, *E. coli* exhibits a different diversification pattern, characterized by lower entropy under chemostat conditions (**Figure 1 G**). This is because the activation of stress-related genes in *E. coli,* monitored here using a *P_ydcS_::GFP* transcriptional reporter as a proxy for RpoS-dependent stress response, does not entail a significant switching cost ^26^. Beyond its analytical capabilities, the Segregostat also enables active control of population dynamics. Indeed, despite the mentioned species-specific differences, it successfully synchronized stress gene expression and reduced overall population heterogeneity in both organisms, compared to chemostat cultures operated at similar dilution rates (**Figure 1 D-I, Supp. Note 1**).

However, all data collected so far has been generated using lab-scale bioreactors. It is well established that scale-up introduces significant challenges, particularly the reduction in global mixing efficiency, which leads to the formation of spatial concentration gradients that can impact cellular physiology. In our context, such gradients are expected to result in zones of glucose excess and starvation, which could limit the effectiveness of the Segregostat technology in regulating the general stress response at the population level by shadowing the environmental changes it is designed to impose. To explore this, we will assess the performance of the Segregostat in two-compartment scale-down bioreactors, which are designed to mimic the heterogeneities encountered in large-scale industrial systems ^7,27^.

### Scaled-down conditions do not lead to an increase of population entropy

To assess the robustness of the Segregostat control under environmental perturbations, we designed a two-compartment scale-down reactor (SDR) consisting of a standard lab-scale stirred-tank bioreactor (STR) connected to a recirculation loop (**Figure 2 A-B**). In this configuration, substrate is typically injected into the recirculation loop. Combined with cellular consumption along the loop, this generates concentration gradients that result in the formation of distinct zones of nutrient excess (red) and starvation (blue) (**Figure 2 A-B**; note the colors are a qualitative illustration, and the local manifestation of excess/starvation conditions may differ between *E. coli* and *S. cerevisiae* due to specific uptake rates). This setup mimics large-scale bioreactor heterogeneities, where subpopulations of cells are stochastically exposed to varying substrate concentrations. As a control, we first cultivated *S. cerevisiae* and *E. coli* in the scale-down reactor (SDR) without applying the Segregostat control procedure (**Figure 2 A-B**). For *E. coli*, two different residence times in the loop (Tr_loop_) of 2 and 8 minutes were tested. Cultivations under these conditions did not result in any noticeable changes in the population profile, as assessed by automated FC, the entropy level being close to the one recorded for lab-scale cultivations (**Figure 2 C-D & Figure 3 C**). In contrast, for *S. cerevisiae*, cultivation in the SDR with a Tr_loop_ of 8 min resulted in a reduction in the overall entropy of the population (**Figures 2 E-F & Figure 3 E**). This outcome can be attributed to the yeast stress response being naturally stimulated by alternating feast-to-famine transitions ^28,29^, mechanistically similar to the strategy employed by the Segregostat (**Figure 1**). In our configuration, cells experience such transitions since they follow a single path through the different conditions, with a stochastic residence time in the stirred tank. Such dynamic environmental conditions have been shown to exert a stabilizing effect on the physiological and proteomic state of cells ^30^. Interestingly, it seems that the residence time in the recirculation loop or the frequencies of the perturbations impact the heterogeneity of the population (**Supp. Note 2**) ^31^. Longer residence times or less frequent perturbations likely promote uniform adaptation and synchronized behavior, whereas shorter residence times lead to more frequent perturbations, eliciting diverse individual responses and increasing population heterogeneity ^31^. In line with this hypothesis, chemostat cultivation of yeast in the SDR with a 4-minute Tr_loop_ did not result in population homogenization (**Figure 3 E**).

**Figure 2:**
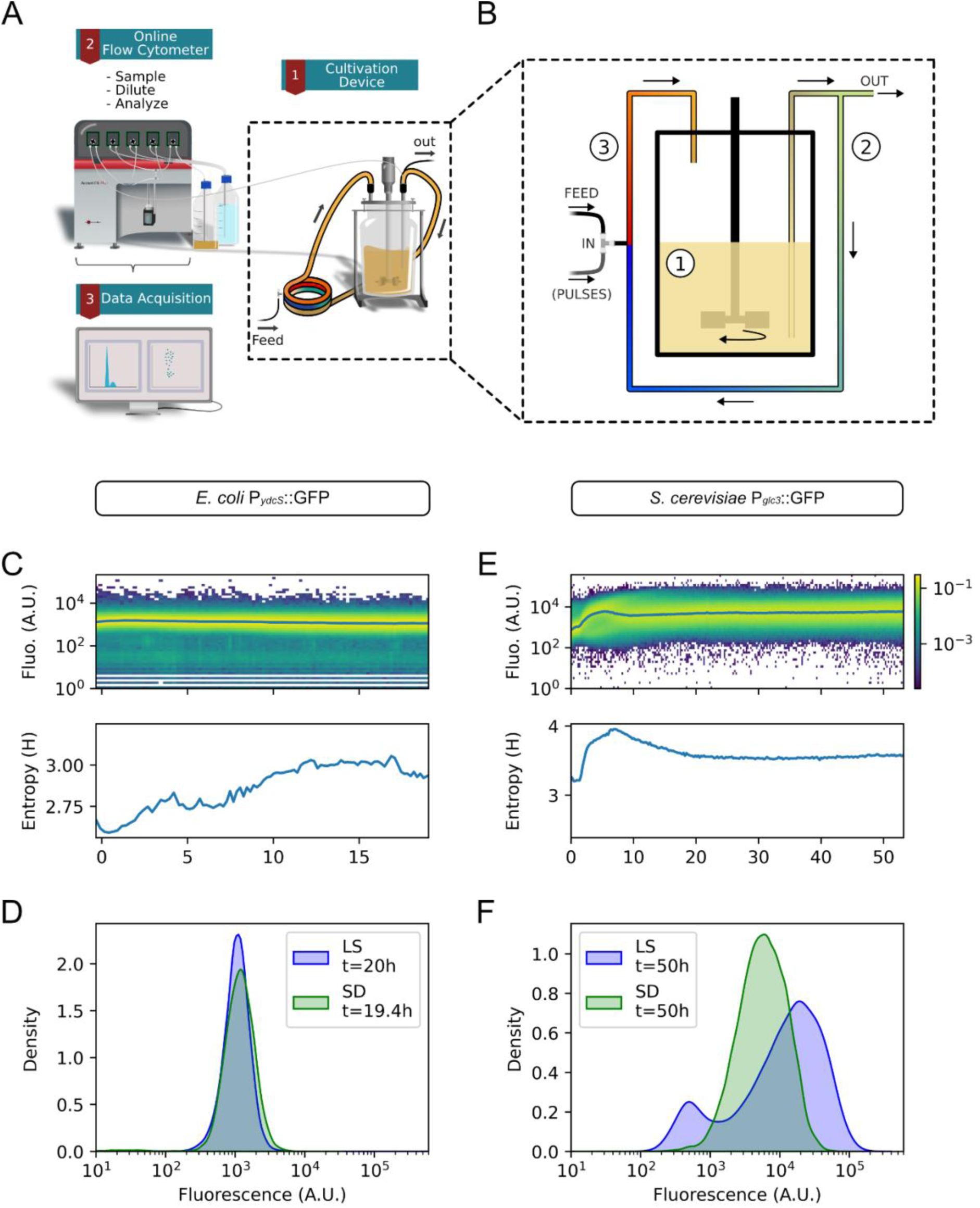
Scale-down effects on population dynamics in the absence of Segregostat regulation. **A** Schematic representation of the online flow cytometry platform for monitoring the scale-down reactor. **B** Simplified Scale-Down design composed by a STR (Zone 1) (V_A_ = 0.8 L) with a recirculation loop (Zone 2 and 3) (V_B+C_ = 0.2 L, V_B_ = 0.067 L, V_C_ = 0.133 L). Feeding, and pulses when present, are performed at 2/3 of the recirculation loop (between zone 2 and 3). **C & E** Fluorescence time-scatter plots and corresponding entropy dynamics from continuous chemostat scale-down cultivation (Tr_loop_ = 8 min) of *E. coli* P*_ydcS_*::GFP (D = 0.3 h⁻¹) (**C**) and *S. cerevisiae* P*_glc3_*::GFP (D = 0.1 h⁻¹) (**E**). **D & F** Comparison of fluorescence distribution between Lab-Scale (LS) and Scale-Down (SD) chemostats conditions at steady state at D=0.3 for *E. coli* (**E**) and D=0.1 for *S. cerevisiae* (**H**). The time indicated corresponds to the duration spent in the condition prior to the measurement.

**Figure 3:**
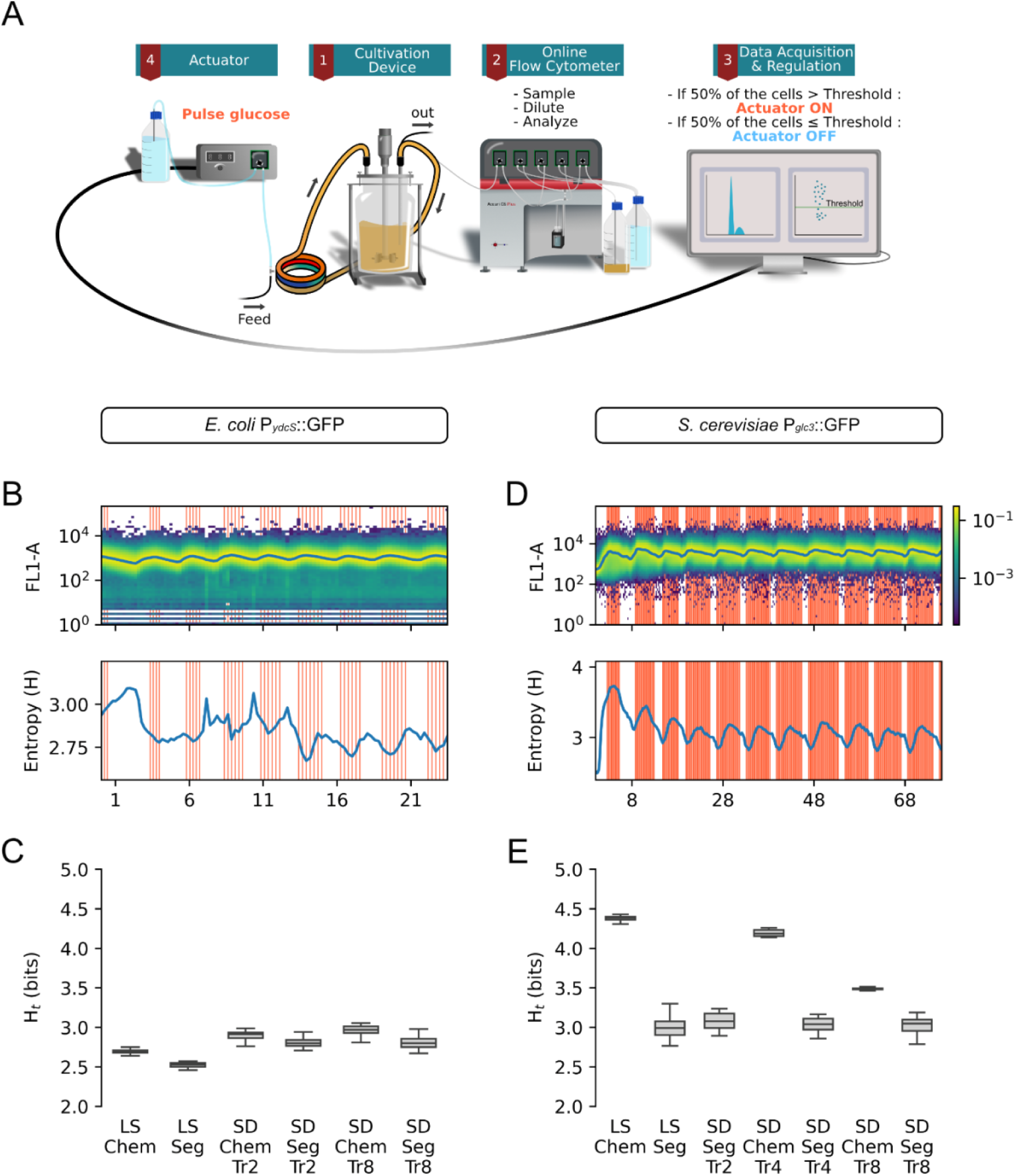
Segregostat exhibits robust control under scale-down conditions. **A** Schematic representation of the Segregostat platform for controlling the scale-down reactor. **B & D** Fluorescence time-scatter plots and corresponding entropy dynamics from continuous Segregostat scale-down cultivation (Tr_loop_ = 8 min) of *E. coli* P*_ydcS_*::GFP (D = 0.3 h⁻¹) (**B**) and *S. cerevisiae* P*_glc3_*::GFP (D = 0.1 h⁻¹) (**D**). **C-E** Basal entropy of *E. coli* (**C**) and *S. cerevisiae* (**E**), calculated from the fluorescence distribution of the population over time, represented as boxplots, for different culture conditions. For all conditions shown in **C–E**, data from the last 12 hours (*E. coli*) or 20 hours (*S. cerevisiae*) were used, corresponding to (quasi-)steady-state under chemostat operation.

These observations highlight the importance of conducting more detailed analyses of cell population behavior under simulated large-scale conditions, notably because, in some cases, environmental heterogeneity can lead to a more homogeneous population-level response.

### Segregostat control leads to robust population profile in scaled-down conditions

We then implemented the Segregostat technology to control gene expression in the scale-down reactor (SDR). To do so, the Segregostat actuator (i.e., glucose pulsing) was applied in the recirculation loop at the same point as the feed input (**Figures 3 A & 2 B**). This setup introduces a key distinction from lab-scale conditions, as glucose pulses now interact with the concentration gradients naturally formed within the SDR. Consequently, the effectiveness of Segregostat-based control is expected to be influenced by these spatial nutrient variations.

Despite this added complexity, Segregostat control consistently led to a reduction in population entropy compared to the corresponding chemostat cultures in the SDR, across all Tr_loop_ tested (**Figure 3 B-E, Supp. Note 3**). This demonstrates that, even under heterogeneous environmental conditions, the Segregostat can successfully control population dynamics. It is worth noting, however, that the reduction in entropy was less pronounced for *E. coli* than for *S. cerevisiae*. This difference reflects the lower heterogeneity of *E. coli* populations in chemostat conditions, due to their smaller switching cost. The environmental forcing imposed by the Segregostat appears to dominate over the perturbations introduced by spatial heterogeneities in the reactor.

Nonetheless, the effectiveness of glucose pulsing in modulating population behavior must be interpreted considering the system’s spatial and temporal dynamics. For instance, monitoring dissolved oxygen levels in the stirred tank during *E.* coli cultivations, both in lab scale bioreactor and in the SDR, revealed that successive glucose pulses likely overlap (**Supp. Note 4**). Specifically, dissolved oxygen levels do not return to their pre-pulse maximum before the next pulse occurs, suggesting that glucose from the previous pulse has not been fully consumed. In addition, for yeast, the delayed rise in oxygen concentration following a series of pulses suggests a metabolic shift toward ethanol consumption. These results highlight how the pulsing strategy influences cellular physiology, a point discussed in greater detail in the following sections.

### Segregostat control induces overflow metabolism and diauxic shifts, but uniformize growth rate distributions

The Segregostat does not abolish scale-down effects; instead, it modulates population physiology by synchronizing the expression of target genes. Unlike the stochastic fluctuations characteristic of large-scale bioreactors, typically mimicked in scale-down reactors, the environmental perturbations imposed by the Segregostat are deliberate and coordinated. In our system, glucose pulsing drives overflow metabolism and diauxic shifts, as reflected in dissolved oxygen traces and base addition profiles (e.g., for both organisms the rate of base addition in SD with Tr = 8 min was 3.5 times higher in Segregostat than in chemostat) (**Supplementary Note 4**). These transitions resemble those observed in spatially heterogeneous industrial fermenters, where their random and unpredictable nature often compromises process efficiency ^6,20^. By contrast, when the timing and magnitude of environmental changes are actively controlled, the detrimental impact of such metabolic shifts can be substantially mitigated. In yeast, for example, glucose pulses induce the Crabtree effect, leading to ethanol excretion; under Segregostat operation, the ethanol produced during the feast phase is fully re-consumed in the subsequent famine phase (**Supplementary Note 4**). This prevents the accumulation of inhibitory by-products and, crucially, reduces the likelihood of a subpopulation entering a persistently stressed state. Such long-term stress responses, often accompanied by growth arrest and metabolic slowdown ^5,14,26^, are far more damaging to culture performance than transient excursions into suboptimal metabolic states.

Beyond stress mitigation, the Segregostat also promotes a more homogeneous population compared to chemostat cultures operated at similar dilution rates (**Figure 3 C & E, Supp. Note 3**). This phenotypic alignment likely results from the strong, population-wide coordination imposed by the pulsing regime, which reduces the emergence of divergent subpopulations despite underlying spatial heterogeneities in the reactor. This feature is particularly relevant for yeast, where the strong trade-off between growth and gene expression leads to pronounced cell-to-cell heterogeneity in both gene expression and growth rate, depending on the environmental conditions and the mode of control (i.e., either simple chemostat or Segregostat) (**Figure 4**). Indeed, homogenous populations, both in terms of stress response and growth rate (**Figure 4B and 4D**), are obtained when the Segregostat is used. This homogenization effect likely arise from the specific environmental perturbation profiles (e.g., in terms of frequency and amplitude) applied in each case. This aspect is further explored in the following section using a simplified cybernetic model.

**Figure 4:**
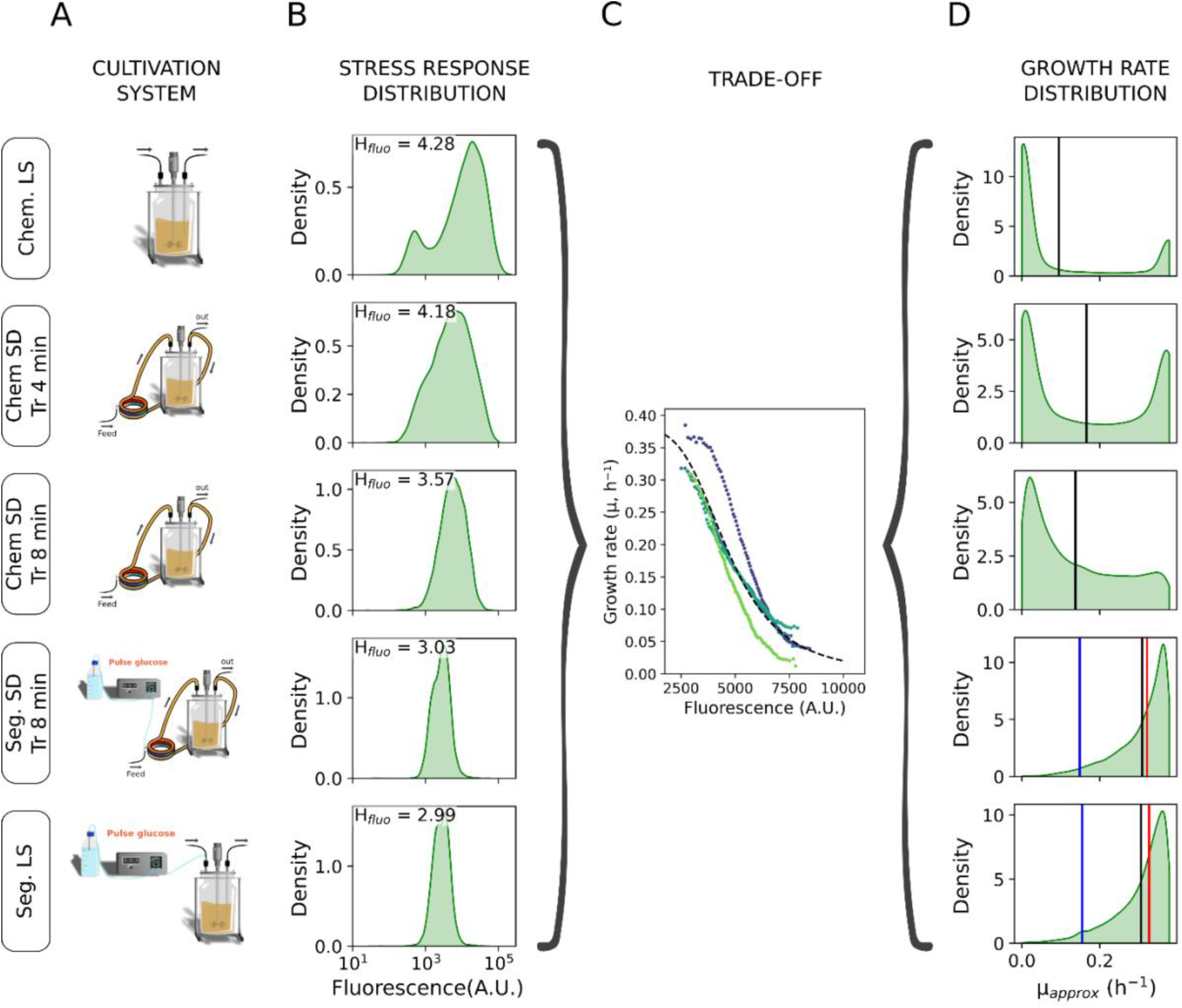
**Impact of bioreactor operating mode on yeast population stress response and growth rate distribution**. **A** Schemes of the different cultivation devices. **B** Distributions of GFP fluorescence related to stress response at steady state. **C** Trade-off curve obtained from Segregostat experiment linking single cell growth rate to the level of stress response. **D** Growth rate distributions inferred from the GFP distribution and the trade-off curve. The vertical black lines mark the mean expected growth rate. Under Segregostat conditions, both the mean growth rate and entropy vary over time; the sample shown was selected because its entropy is close to the median across all samples from the corresponding Segregostat experiment. Blue and red vertical lines indicate the minimum and maximum expected growth rates, respectively.

### Segregostat-imposed environmental perturbations dominate over mixing-induced heterogeneities and drive population entrainment

In large-scale bioreactors, cells experience irregular perturbations of relatively high frequency and low amplitude due to mixing imperfections (**Figure 5 A**, **Supp. Note 6**) ^7,32^. Such dynamic heterogeneity in glucose concentration was mimicked in our scale-down reactor (**Figure 5 C**). In contrast, the Segregostat, through real-time monitoring and control of stress expression, generates feast-to-famine transitions (by alternating between pulse series and pulse-free phases) that align with host-circuit dynamics (**Figure 5 B & D**). These regular perturbations can be viewed as a quasi-square-wave signal characterized by relatively low frequency and high amplitude (**Figure 5 D**). The step time of this signal, and thus the duration of each phase (feast, when glucose concentration is not limiting, and famine, when cells experience stress), is automatically determined by the Segregostat, allowing the population to respond homogeneously (**Figure 3 D & E**).

**Figure 5:**
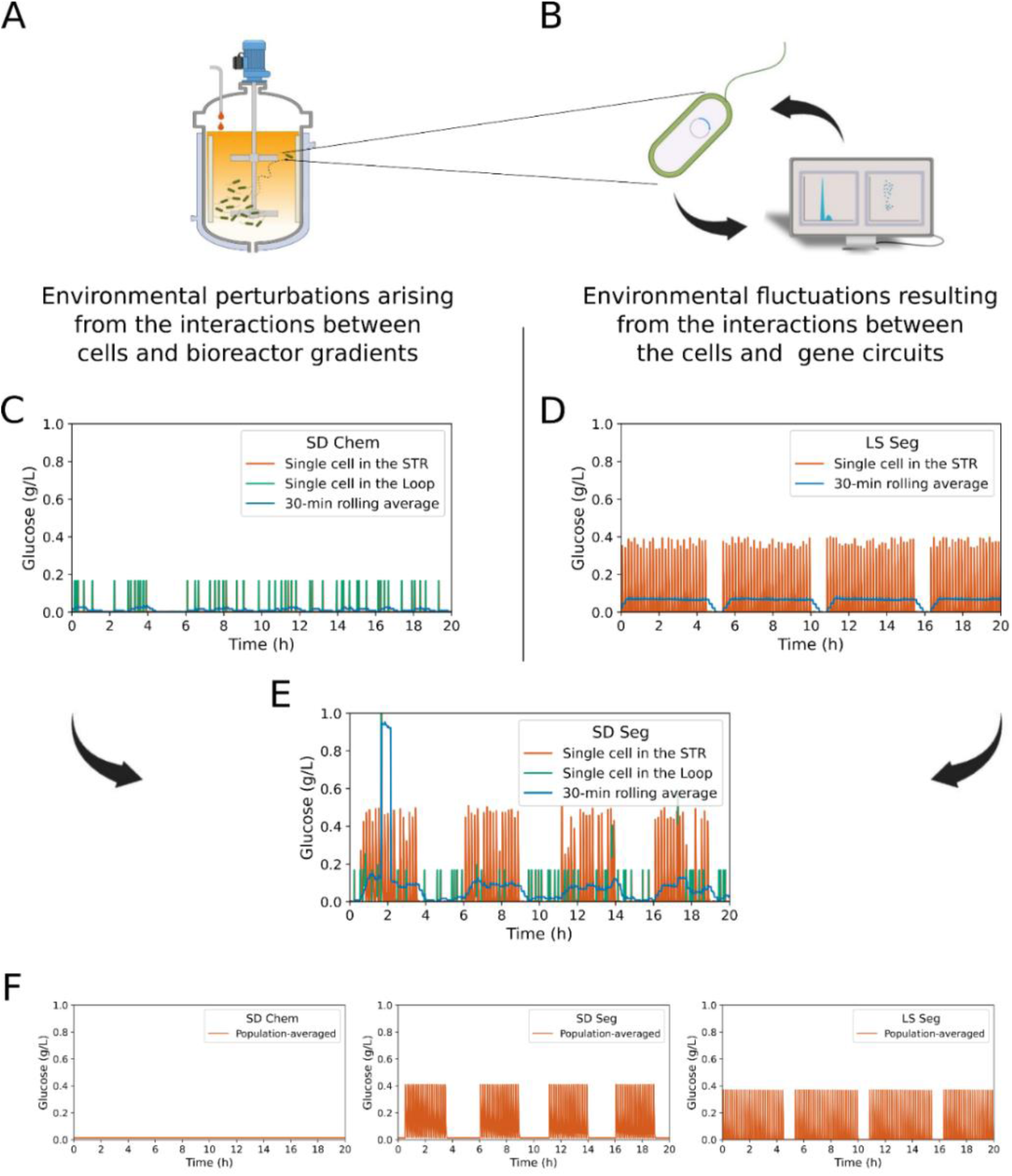
Different types of interactions drive different kinds of environmental fluctuations. **A** The interactions between cells and bioreactors lead to stochastic cell lifeline, characterized by low amplitudes and high frequencies. **B** The interactions between the gene circuit and the cellular host, enforced by the Segregostat, leads to environmental fluctuations with high amplitude and low frequency. **C-E** Simulated single-cell lifelines in terms of glucose concentrations perceived in continuous cultures. Lines are orange when the cell is in the STR and green when it is in the recirculation loop. Blue lines indicate a 30-min rolling average. (**C**) In SD conditions (Tr_loop_ = 4min) without control, where interactions between cells and bioreactors dominate, (**D**) In LS Segregostat conditions, where fluctuations following the interactions between the gene circuit and the cellular host dominate, and (**E**) in SD Segregostat (Tr_loop_ = 4min) were both interactions are present. **F** The corresponding mean glucose concentration perceived by approx. 1600 cells over time, showing that regular perturbations of the Segregostat dominate even in scale-down conditions.

When both effects are superimposed (**Figure 5 E**), Segregostat-imposed fluctuations dominate, regardless of the residence time in the loop, and the quasi-square-wave signal is robustly transmitted to the cells. In other words, SD-induced environmental perturbations do not disrupt the Segregostat signal. Analysis of the population-averaged glucose concentrations perceived over time further highlights that the Segregostat perturbations are coherently perceived by all the cells, whereas SD perturbations are randomly perceived in the loop (**Figure 5 C-F**). In this sense, the Segregostat can be regarded as a "noise-canceller" for the cells.

These observations are consistent with well-established principles in systems and synthetic biology. Both modeling ^33–35^ and experimental ^36–38^ studies have shown that cellular systems evolve two modes of phenotypic switching depending on the nature of environmental perturbations to enhance population fitness. When fluctuations are irregular in amplitude and timing and occur at high frequency, stochastic switching between gene circuit states is favored, thereby increasing cell-to-cell heterogeneity. Conversely, when fluctuations are more regular, deterministic switching strategies prevail, leading to more homogeneous populations. Our data align with these predictions (**Figure 4** & **Figure 6**): population entropy is higher when perturbations from cell–environment interactions dominate (as in large-scale bioreactors) (**Figure 5 A** & **Figure 3 E**), while entropy is lower under Segregostat control, where fluctuations are primarily governed by cell–gene circuit interactions (**Figure 5 B** & **Figure 3 E**).

**Figure 6:**
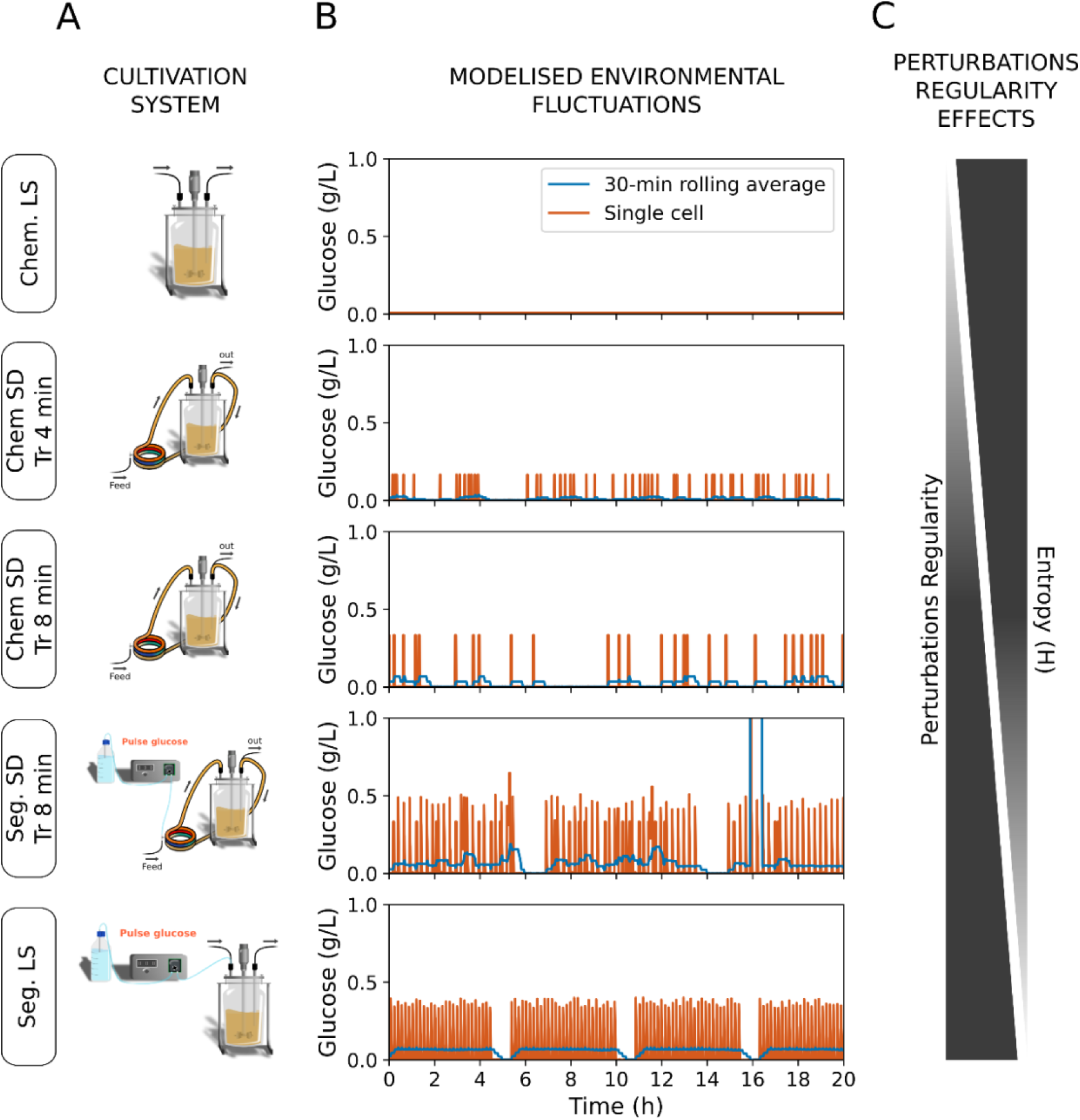
**Impact of bioreactor operating mode and the corresponding environmental perturbations profiles on yeast population heterogeneity**. **A** Schemes of the different cultivation devices. **B** Simulated single-cell lifelines in terms of glucose concentrations perceived in corresponding culture conditions. **C** Consistency of environmental perturbations (i.e., suitable frequencies, amplitudes and step-times) and heterogeneity of the population in corresponding culture conditions. In this representation, we considered the perturbations arising in a chemostat at lab-scale as extremely frequent and of very small amplitude.

Interestingly, the homogeneization of the population observed for longer residence times in the loop (Tr_Loop_ = 8min) could be explained by the same principles. With a longer residence time, cells are exposed to higher glucose concentrations for extended periods (**Figure 6**). Although these fluctuations remain random, their amplitude, frequency, and duration allow the cells to respond in a more similar manner, as discussed above.

## Discussion

Controlling cell population dynamics and phenotypic diversification is a central challenge in systems and synthetic biology, where engineered gene circuits are expected to elicit uniform responses upon stimulation ^39^. Traditionally, this problem has been addressed through a variety of control strategies, most notably cell-machine interfaces. Such interfaces modulate complex host-circuit interactions by triggering feedback, for instance based on the expression level of a target gene, thereby promoting more homogeneous gene expression across single cells within a population ^10,40^.

In this work, we introduce an additional component by considering environmental perturbations, such as those typically encountered by cell populations in heterogeneous bioreactor environments ^6^. Specifically, we focused on the control of the general stress response in two industrially relevant hosts, *Saccharomyces cerevisiae* and *Escherichia coli*, and performed scale-down cultivation experiments to evaluate the control performance of a previously developed cell-machine interface known as the Segregostat ^5,13^. This system employs automated flow cytometry to continuously monitor cellular physiology in real time and autonomously initiates feast-to-famine transitions, thereby coordinating stress responses across the population.

Remarkably, the Segregostat demonstrated robust control performance for all scale-down conditions tested. This robustness likely arises because the Segregostat imposes and amplifies perturbations that align with host-circuit dynamics and that are not disrupted by those introduced by bioreactor constraints, thereby preserving control performance. By directly monitoring gene circuit activation, the Segregostat captures intrinsic dynamics ^41^ that, like other components of cellular regulatory networks, have evolved under selective pressures imposed by environmental fluctuations and their timing ^33,36,42,43^. As a result, the environmental perturbations imposed by the Segregostat synchronize with the intrinsic rhythms emerging from the interplay between the cellular system and its stress-responsive gene circuit. These homogeneous, periodic, and precisely defined fluctuations promote responsive switching and coordinated gene expression across the population. This entails a reduction in entropy, more pronounced in *S. cerevisiae* than in *E. coli*, reflecting the lower switching cost and associated baseline heterogeneity of *E. coli* populations.

Dynamic environmental conditions have been reported to stabilize gene expression ^31^, the physiological and proteomic states of cells ^30^, and even the structure of microbial communities ^44–46^. In some cases, such conditions were predicted by models, while in others they arose unintentionally from uncontrolled fluctuations rather than deliberate control. We also observed a comparable homogenizing effect in chemostat experiments under scale-down conditions with extended residence times in the recirculation loop. Importantly, the temporal features of these fluctuations are a critical determinant of the outcome ^31,47^. This raises the question of whether fixed perturbations, applied at frequencies matching the time constants of gene expression, could replicate the effects achieved with Segregostat control. Crucially, however, the Segregostat goes beyond such chance occurrences: its closed-loop control autonomously identifies the relevant time constants and imposes fluctuations precisely matched to host–circuit dynamics, thereby ensuring robust population-level control.

The control strategy applied in this study was intentionally simple, relying on a single threshold for one monitored gene, but the framework is readily extensible. For example, cell size has already been used as a control variable to discriminate between yeast and bacteria in co-culture systems ^48^. A natural next step would be to couple this approach to circuits directly linked to a product of interest, or to synthetic gene networks engineered for bioproduction. Incorporating multi-reporter strains would further allow simultaneous monitoring of multiple phenotypic traits, paving the way for more sophisticated, multi-parametric control strategies ^31^. Ultimately, the pulse sequences identified by the Segregostat in scale-down conditions could be transferred in open-loop mode to large-scale bioreactors and validated under industrially relevant conditions.

In summary, the Segregostat emerges as a promising tool to stabilize long-term cultures by ensuring an optimal trade-off between growth and stress response. In the future, this principle could also be applied to bioproduction system by finding the optimal trade-off between growth and product formation, thereby preventing the selection of less productive phenotypes ^30, 49^. By aligning environmental perturbations with the intrinsic dynamics of regulatory networks, this strategy offers a powerful means to achieve robust, coordinated gene expression at scale. Looking forward, we believe that future bioprocess design should adopt a more cell-centric perspective: beyond optimizing physical parameters of the reactor, such as mixing or aeration, it is crucial to integrate the physiology and regulatory logic of the host as central elements of the system. Cells should be regarded not merely as passive entities subjected to environmental perturbations, but as active partners whose intrinsic dynamics can be harnessed for robust and productive processes. Our work contributes to building this cell-centric mindset by strengthening the integration between systems biology, synthetic biology, and bioprocess engineering, ultimately fostering the design of next-generation bioprocesses that are both efficient and resilient.

## Material and methods

### Code and data availability

Scripts used for data analysis, figure generation, and for the cybernetic model are available at https://gitlab.uliege.be/mipi/published-software/2025-scaledownsegregostat. The raw data have been deposited in Zenodo under DOI: https://doi.org/10.5281/zenodo.17453611.

### Microbial strains and growth media

The bacterial model used was the *Escherichia coli* MG1655 P*ydcS*::GFPmut2 strain from the Zaslaver collection ^50^. This strain carries a plasmid containing a transcriptional reporter for P*_ydcS_* and a kanamycin resistance marker. The yeast used in this work was the *Saccharomyces cerevisiae* CEN.PK 113-7D strain with the chromosomal integration of a reporter cassette P*_glc3_*::eGFP ^51,52^. The rationale for selecting P*_ydcS_* and P*_glc3_* as reporters of the general stress response in *E. coli* and *S. cerevisiae* has been described previously ^26^.

Unless otherwise specified, the following media were used. *E. coli* has been cultivated in a mineral medium containing (in g/l), K_2_HPO_4_: 14.6, NaH_2_PO_4_ ⋅ 2H_2_O: 3.6, Na_2_SO_4_: 2, (NH_4_)_2_SO_4_: 2.47, NH_4_Cl: 0.5, (NH_4_)_2_−H−citrate: 1, glucose: 5, thiamine: 0.01. The medium is supplemented with a trace element solution added at a ratio of 11 ml/l and assembled from the following solutions (in g/l), 3/11 of FeCl_3_ ⋅ 6H_2_O: 16.7, 3/11 of EDTA: 20.1, 2/11 of MgSO_4_: 120, and 3/11 of a metallic trace element solution. The metallic trace element solution contains (in g/L): CaCl_2_ ⋅ 2H_2_O: 0.74, ZnSO_4_ ⋅ 7H_2_O: 0.18, MnSO_4_ ⋅ H_2_O: 0.1, CuSO_4_ ⋅ 5H_2_O: 0.1, CoSO_4_ ⋅ 7H_2_O: 0.21. For plasmid maintenance, it has been supplemented with kanamycin (50 mg/l). The trace element solution, the thiamine and the antibiotic were filter-sterilized (0.22 µm) before supplementation of the other component that were heat-sterilized (at 120°C).

*S. cerevisiae* has been cultivated in a mineral medium based on the recipe of Verduyn et al., (1992). It contains, per liter, (NH_4_)_2_SO_4_: 5 g, KH_2_PO_4_: 3 g, MgSO_4_ ⋅ 7H_2_O: 0.5 g, EDTA: 95.55 mg, ZnSO4 · 7H2O: 22.5 mg, MnCl2 · 4H2O: 5 mg, CoCl2 · 6H2O: 1.5 mg, CuSO4 · 5H2O: 1.5 mg, Na_2_MoO_4_ · 2H_2_O: 2 mg, CaCl_2_ · 2H_2_O: 22.5 mg, FeSO_4_ · 7H_2_O: 15 mg, H_3_BO_3_: 5 mg, KI: 0.5 mg and glucose: 7.5 g. After heat sterilization (at 120°C) of those components, the filter-sterilized (0.22 µm) vitamins were added. The final medium contains per liter, D-biotin: 0.05 mg, calcium pantothenate: 1 mg, nicotinic acid: 1 mg, myo-inositol: 25 mg, thiamine HCl: 1 mg, pyridoxine HCl: 1 mg and para-aminobenzoic acid: 0.2 mg. For bioreactor cultivations, 100 µL of TEGO Antifoam KS 911 were added per liter of medium.

### Continuous cultivations operations

The operating parameters of continuous cultures are provided in the **Supp. Tables**.

To prevent unwanted reporter activation, strains were cultivated in two successive precultures. Both steps were carried out in 500 ml or 1 L flasks containing 50 or 100 ml of medium, respectively, incubated at 37 °C for *E. coli* or 30 °C for *S. cerevisiae*, with agitation at 150 rpm. The first preculture was started either from a glycerol stock or from a single colony grown on agar plate (LB supplemented with 50 mg/L kanamycin for *E. coli*, YPD for *S. cerevisiae*) and incubated overnight. The second preculture was inoculated with this culture to reach an initial OD_600_ ≈ 0.5 and incubated until sufficient biomass was obtained for bioreactor inoculation (approximately 3 h for *E. coli* and 6 h for *S. cerevisiae*). Batch phases were initiated at an OD_600_ of about 0.1 (or 0.2 for experiments performed in the Biostat B-Twin bioreactor).

Continuous cultivations were conducted in either Biostat B-Twin (Sartorius, Göttingen, Allemagne) or in Bionet F1 bioreactor (Bionet, Murcia, Spain), with a working volume of 1 L, an agitation speed of 1200 rpm for *E. coli* or 1000 rpm for *S. cerevisiae*, and an aeration rate of 1 VVM. Temperature was maintained at 37 °C for *E. coli* and 30 °C for *S. cerevisiae*, while pH was regulated at 7 and 5, respectively. Each cultivation began with a batch phase that continued until the carbon source was fully depleted (as indicated by a rise in dissolved oxygen) or, for yeast, after approximately 15 h. Chemostat conditions were applied at D = 0.3 h⁻¹ for *E. coli* and D = 0.1 h⁻¹ for *S. cerevisiae*, with each condition maintained for at least five residence times. In Segregostat experiments, a feedback control algorithm (custom MATLAB script based on online flow cytometry data, see below) actuated a pump to deliver glucose pulses. Because of these pulses, the effective dilution rate temporarily exceeded the nominal chemostat values (up to 0.33 h⁻¹ for *E. coli* and 0.125 h⁻¹ for *S. cerevisiae*). Pulses were triggered when more than 50% of the population exceeded a predefined fluorescence threshold.

The simplified scale-down bioreactor consisted of a stirred-tank reactor (working volume 0.8 L) coupled with a recirculation loop (working volume 0.2 L) (**Figure 2B**). The recirculation loop was composed of a 10 m long, 0.5 cm diameter silicone tube. Feeding and pulse additions were applied at two-thirds of the tube length i.e., one-third before returning to the stirred tank. Chemostat and Segregostat cultures were performed at different residence times in the recirculation loop (see **Supp. Tables** for the list of tested conditions). All other operating parameters were identical to those used in the lab-scale bioreactors described above. No limitations other than the carbon source were observed (**Supp. Note 5**).

### Continuous cultivations monitoring and data treatment

Dissolved oxygen and pH were monitored in all experiments. For experiments performed in Bionet F1 bioreactor, biomass was monitored with Hamilton Dencytee RS485 probes. Single-cell dynamics were monitored throughout the cultures using an automated flow cytometry setup. This system combined a custom-built sampling device (integrated in the Segregostat platform (Henrion et al., 2023a; Sassi et al., 2019)) with a benchtop flow cytometer (BD Accuri C6 or BD Accuri C6+). Every 12 minutes, a sample was withdrawn from the bioreactor, diluted in PBS, and subsequently analyzed.

For measurements obtained with the BD Accuri C6, the cytometer was operated with custom fluidics setting (24 µL/min, 8 µm core), and the FSC-H threshold was adjusted to 20,000 for *E. coli* and 80,000 for *S. cerevisiae*. When using the BD Accuri C6+, the flow rate was set to “medium” (35 µL/min, 16 µm core), and the threshold was set to 15,000 for *E. coli* and 80,000 for *S. cerevisiae*. Under these configurations, roughly 40,000 cells were acquired per sample. GFP fluorescence was recorded in the FL1-A channel (excitation at 488 nm and emission filter 533/30 nm).

To enable direct comparison between replicates measured on different flow cytometers, data from the BD Accuri C6 were transformed as previously described (Delvenne et al., 2025). Flow cytometry datasets were pre-processed by discarding zero values and doublets. Entropy (H) was calculated using the formula below, where m is the number of bins and *pᵢ* the probability of a data point falling into bin *i*. In our case, the fluorescence distribution was discretized into 50 logarithmically spaced bins spanning from 1 to 10⁶ a.u.

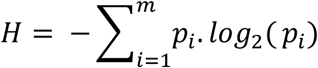

### Cybernetic modelling of continuous cultivations

Models for the different continuous cultivations, with and without plug flow reactor (PFR) loops, were performed using a simplified version of a toolbox published in a previous work (MONKS) ^46^. The toolbox and data files used for the calculation of each model as well as the jupyter notebooks with each of the performed models can be found in the GitLab repository referenced above.

The main difference are the modifications to adapt a PFR to the main MONKS modeling toolbox. To this end, the model was set on creating volumetric cells with a fraction of the total volume of the CSTR and PFR volumes. Initial Biomass is then divided into each volumetric cell. These volumetric cells then virtually exist inside the CSTR until randomly selected to enter the PFR. The selection is performed every delta time and made with a uniform probability density function for the remaining volumetric cells inside the CSRT. The number of volumetric cells selected to enter the PFR is calculated considering the delta volume equal to the fraction of the volume needed to enter PFR according to the flow rate for each experiment (calculated from the retention time in the PFR and the total volume of the PFR). Once in the PFR the volumetric cells pass through it as sections of volumes and only updated in their concentrations until arriving the section of the feed and pulse ports (2/3 of the total PFR volume). Feed is continuously updated at this point, and pulses are performed according to the experimental files. Once cells exit the PFR the remaining mass of the feeds and pulses are used to update the concentration in all the cells of the CSTR. For all simulations between 1500 and 2000 volumetric cells were used to simulate different cell experimental lifelines as each volumetric cell contains the biomass information and the environmental substrates and products available, at this point and for each delta time MONKS cybernetic modeling approach is used to calculate the biomass growth, substrate consumption and product formation within each volumetric cell. Finally, in the CSTR the effect of each cell is distributed for all cells as environmental information, while in the PFR the environmental information is not distributed and therefore the biomass can only use substrate directly in their own volumetric cell.

## Acknowledgement

This research was supported by ERA CoBioTech/EU H2020 project (grant 722361) ‘ComRaDes’, a public–private partnership between the University of Stuttgart, TU Delft, University of Liège, DSM, Centrient Pharmaceuticals, and Syngulon. FD, MD and JMA are very grateful to Vincent Vandenbroucke, Samuel Telek and Andrew Zicler for the help for calibrating and setting the bioreactor experiments.

