## Supplementary material for "Overriding bioprocess perturbations with a cell–machine interface for reliable microbial stress-response control"

Supplementary materials for

### A cell-machine interface enables robust control of stress response in microbial populations upon bioprocess scale-up/down

First version

Mathéo Delvenne[a], Juan Andres Martinez[a], Cees Haringa[b], Henk Noorman[b,c], Steven Minden[d], Ralf Takors[e], and Frank Delvigne[a,1]

[a] Terra Research and Teaching Centre, Microbial Processes and Interactions (MiPI), Gembloux Agro-Bio Tech, University of Liège, Gembloux, Belgium
[b] Department of Biotechnology, Delft University of Technology, Van der Maasweg 9, 2629 HZ Delft, the Netherlands
[c] dsm-firmenich, A. Fleminglaan 1, 2613 AX Delft, the Netherlands
[d] Institute for Biological Interfaces 5 (IBG-5), Biotechnology and Microbial Genetics, Karlsruhe Institute of Technology (KIT), Hermann-von-Helmholtz-Platz 1, 76344 Eggenstein-Leopoldshafen, Germany
[e] Institute of Biochemical Engineering, University of Stuttgart, Allmandring31, 70569 Stuttgart, Germany

19th November 2025

### Contents

**Supplementary Note 1.**
**Distribution of stress response activation in *S. cerevisiae* at different dilution rates** **2**

**Supplementary Note 2.**
**Impact of pulsed glucose feeding on the population heterogeneity** **3**

2.1. Extended materials and methods . . . 3

**Supplementary Note 3.**
**Basal entropy values for all experiments and their replicates** **4**

**Supplementary Note 4.**
**Examples of monitored reactor parameters indicating overflow metabolism** **5**

4.1. Extended materials and methods . . . 6

**Supplementary Note 5.**
**Absence of oxygen limitation in the recirculation loop** **7**

5.1. Extended materials and methods . . . 8

**Supplementary Note 6.**
**CFD simulation of industrial aerated fermentation of *S. cerevisiae*** **9**

**Tables** **10**

7.1. Bioreactors cultivations . . . 10

**Data and Code accessibility** **13**

8.1. Data . . . 13
8.2. Code . . . 13

**Supplementary References** **14**

### Supplementary Note 1. Distribution of stress response activation in *S. cerevisiae* at different dilution rates

In changestat continuous cultures, the feed pump speed was either increased (accelerostat, A-stat) or decreased (decelerostat, D-stat) in steps of 1 rpm every two hours, resulting in a dilution rate change of 0.004 $h^{-1}$ per step [1]. This strategy was chosen to ensure a quasi-steady state at each step.

Chemostat cultures of *S. cerevisiae* were performed at D = 0.1 $h^{-1}$. However, with the addition of pulses in Segregostat, the dilution rate was slightly increased. We therefore compared the entropy values of Segregostat cultures with those of changestat cultures at similar dilution rates (Figure S1).

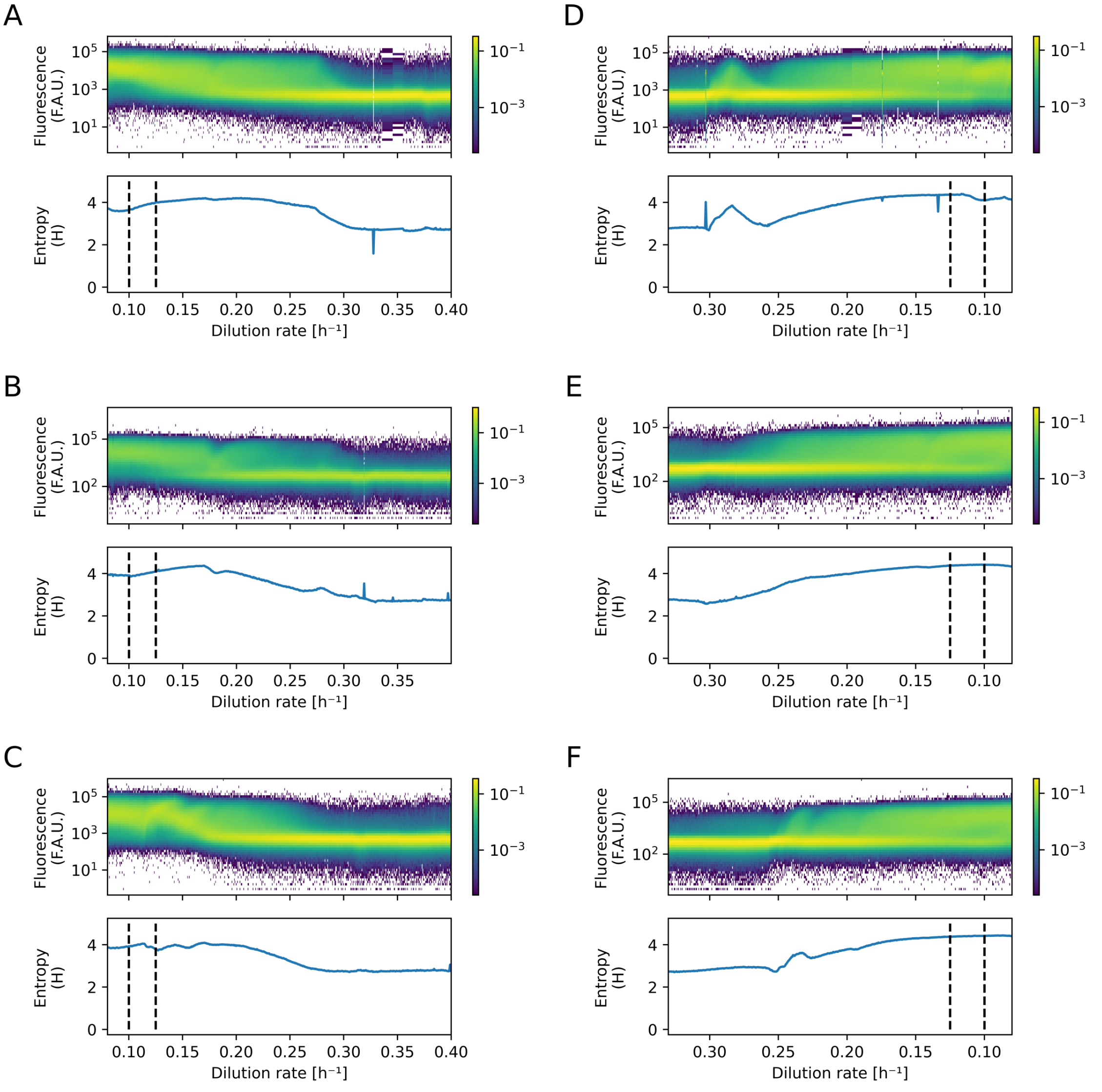


Figure S1: Scatter plots of single-cell fluorescence of *S. cerevisiae* as a function of dilution rate and the corresponding entropy values, for accelerostats (A-C) and decelerostats (D-F) in triplicates. Black dotted vertical lines indicate the minimum and maximum dilution rates applied in Segregostat cultures of *S. cerevisiae*. Data from Maximilian Sehrt's work [1].

### Supplementary Note 2. Impact of pulsed glucose feeding on the population heterogeneity

When glucose was supplied in pulses rather than continuously, lower pulse frequencies resulted in a more homogeneous population (Figure S2).

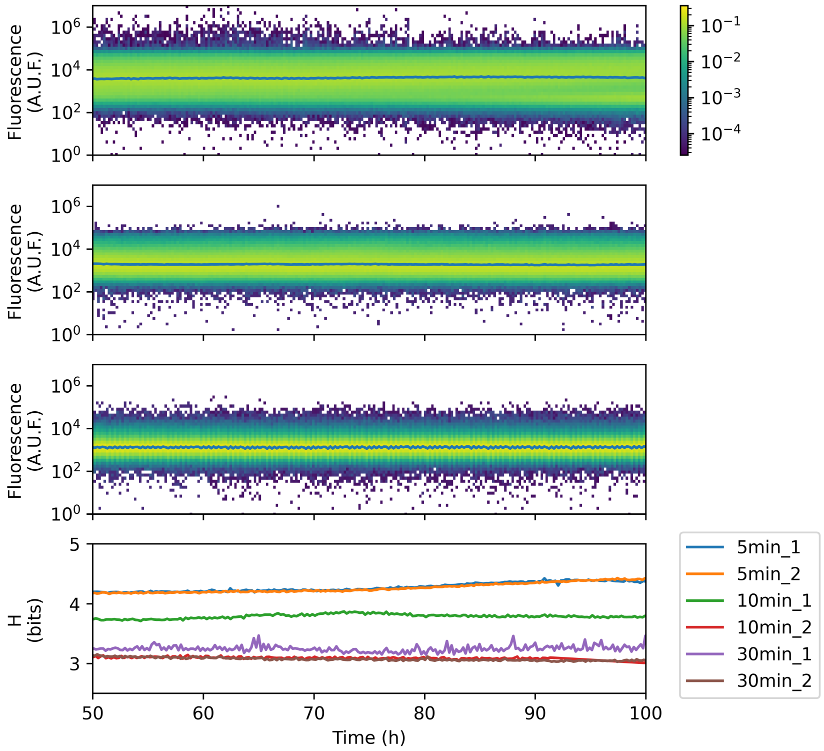


Figure S2: Population heterogeneity of *S. cerevisiae* under pulsed glucose feeding at different frequencies. Single-cell fluorescence distributions are shown for pulsed feeding every 5 min (top), 10 min (middle), and 30 min (bottom). The lower panel shows the corresponding Shannon entropy values (H) over time for two biological replicates per condition. Lower pulse frequencies (e.g. 30 min) result in reduced entropy values, indicating more homogeneous populations.

#### 2.1. Extended materials and methods

Continuous cultures with pulsed glucose feeding were performed in a Biostat B-Twin bioreactor (Sartorius, Göttingen, Germany), in the same way as the chemostat cultures at $D = 0.1\ h^{-1}$ (see Materials and Methods of the main document), except for the feeding strategy. In this case, medium without a carbon source was continuously supplied, while glucose was provided in pulses (from a 20% solution, delivered every 5, 10, or 30 min). The total amount of glucose supplied over time was the same as in the chemostat cultivation at $D = 0.1\ h^{-1}$.

### Supplementary Note 3. Basal entropy values for all experiments and their replicates

Basal entropy values for all replicates are shown in Figure S3 for *E. coli* and in Figure S4 for *S. cerevisiae*.

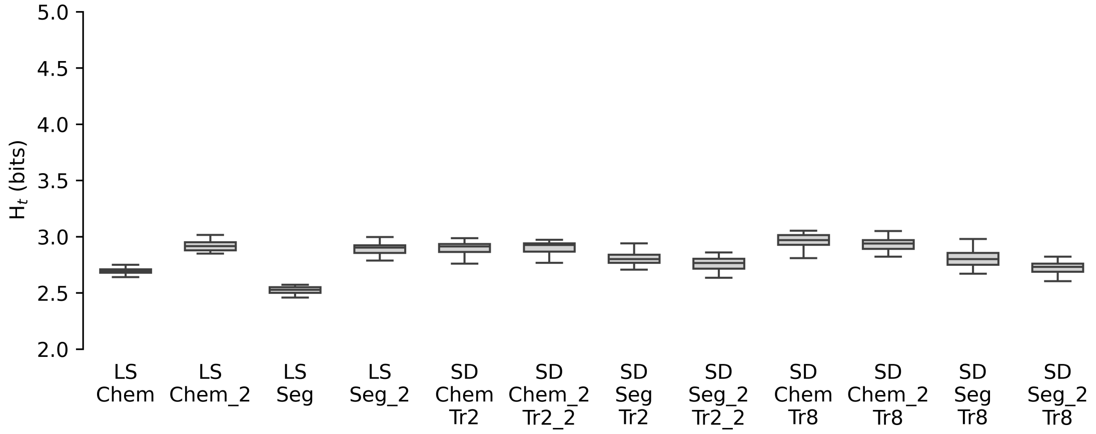


Figure S3: Basal entropy of *E. coli*, calculated from the fluorescence distribution of the population over time, represented as boxplots, for different culture conditions. For all conditions shown, data from the last 12 hours of the condition were used, corresponding to (quasi-)steady-state under chemostat operation.

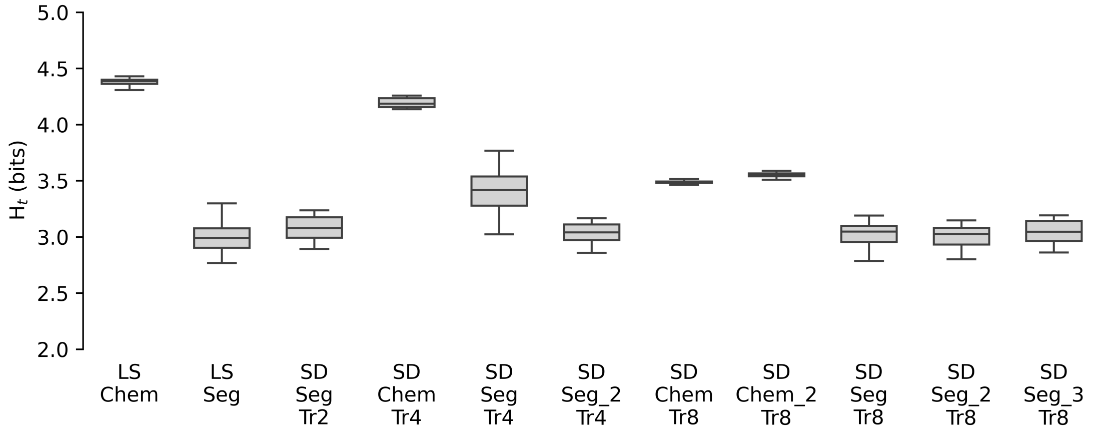


Figure S4: Basal entropy of *S. cerevisiae*, calculated from the fluorescence distribution of the population over time, represented as boxplots, for different culture conditions. For all conditions shown, data from the last 20 hours of the condition were used, corresponding to (quasi-)steady-state under chemostat operation.

### Supplementary Note 4. Examples of monitored reactor parameters indicating overflow metabolism

Differences in pH and dissolved oxygen between chemostat and segregostat cultures indicate overflow metabolism in the latter cultivation mode. Furthermore, in experiments where pH control was achieved solely through base addition (without acid supplementation), the base addition rate reflects the acidification of the medium. These variables are shown in Figure S5, and approximated base addition rates are presented in Figure S6.

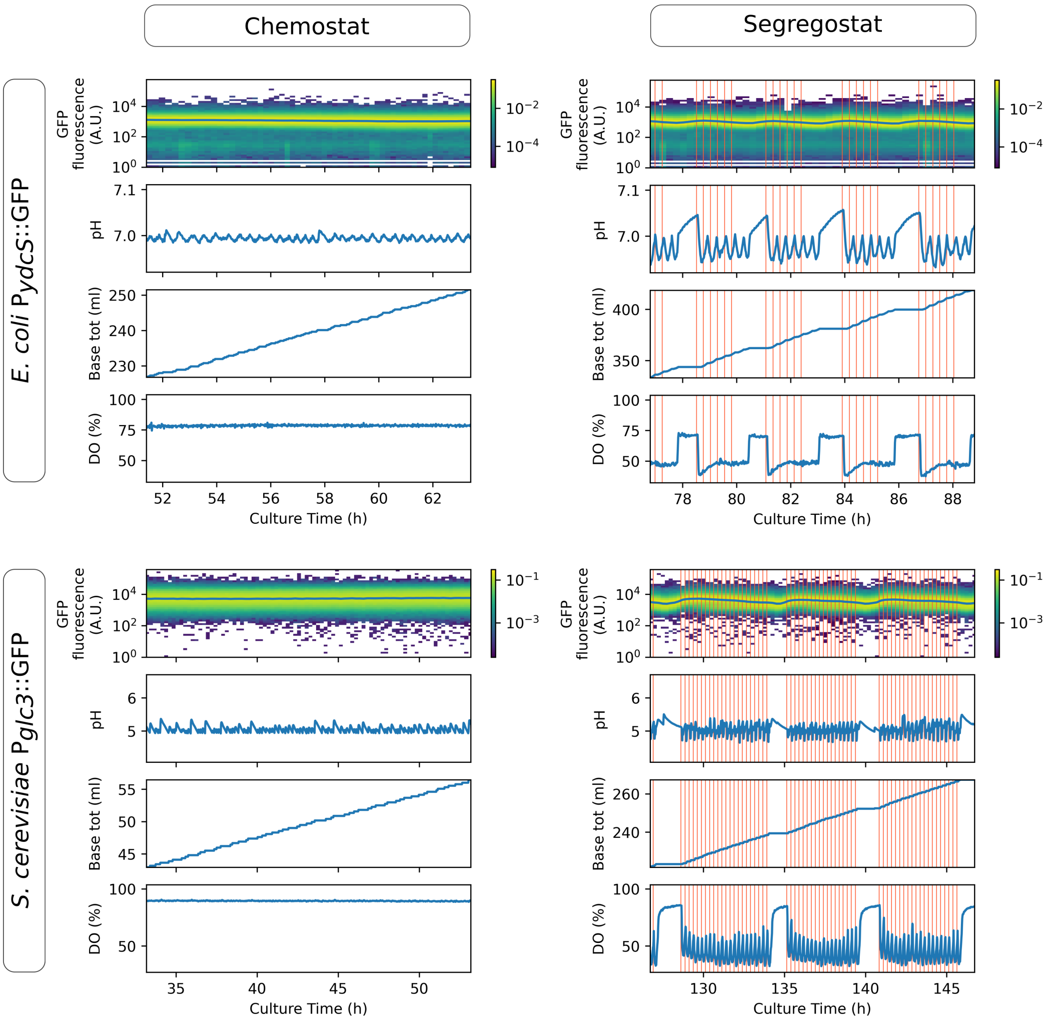


Figure S5: Examples of monitored reactor parameters indicating overflow metabolism in representative cultures of *E. coli* and *S. cerevisiae* under chemostat and Segregostat scale-down (Tr = 8 min) conditions. For each culture, single-cell GFP fluorescence distributions, pH profiles, total base addition, and dissolved oxygen (DO) are shown over time. Red vertical lines in segregostat experiments indicate the timing of glucose pulses. Segregostat cultures exhibit higher base addition rates, stronger pH fluctuations, and marked oscillations in DO compared to chemostats, reflecting overflow metabolism induced by glucose pulses. For instance, in *S. cerevisiae* Segregostat cultures, the dissolved oxygen (DO) level does not return to its maximal value after a glucose pulse, suggesting that the following pulse occurs before carbon source availability becomes limiting. Even after the final pulse of a series, DO requires a delay before stabilizing, which suggests a subsequent re-consumption of ethanol.

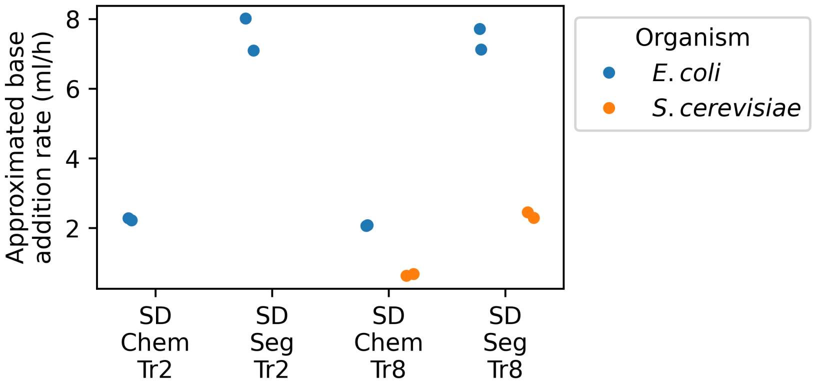


Figure S6: Approximated base addition rates in different culture conditions. Scatter plot of the average base addition rates (ml/h) in cultures of *E. coli* (blue) and *S. cerevisiae* (orange) under scale-down (SD) chemostat and Segregostat experiments with different retention times in the recirculation loop (2 and 8 min). Each point corresponds to an individual replicate.

#### 4.1. Extended materials and methods

The base addition rate was approximated from the difference in the cumulative base added over a fixed time window taken at the end of each culture. For *E. coli*, a 12 h window was used, whereas for *S. cerevisiae* a 20 h window was considered. This approximation is only valid for experiments in which pH control was performed exclusively through base addition, and not through combined acid and base titration.

### Supplementary Note 5. Absence of oxygen limitation in the recirculation loop

The recirculation loop consisted of silicone tubing, which is permeable to oxygen. In addition, small air bubbles were entrained into the loop together with the culture broth. Oxygen limitation was therefore not expected. This assumption was verified by measuring dissolved oxygen concentrations at different positions along the recirculation loop. An example of these measurements is shown in Figure S7.

It is worth noting that small temperature variations may have occurred in the recirculation loop, since this part of the system was not temperature-controlled; temperature regulation was applied only in the stirred tank.

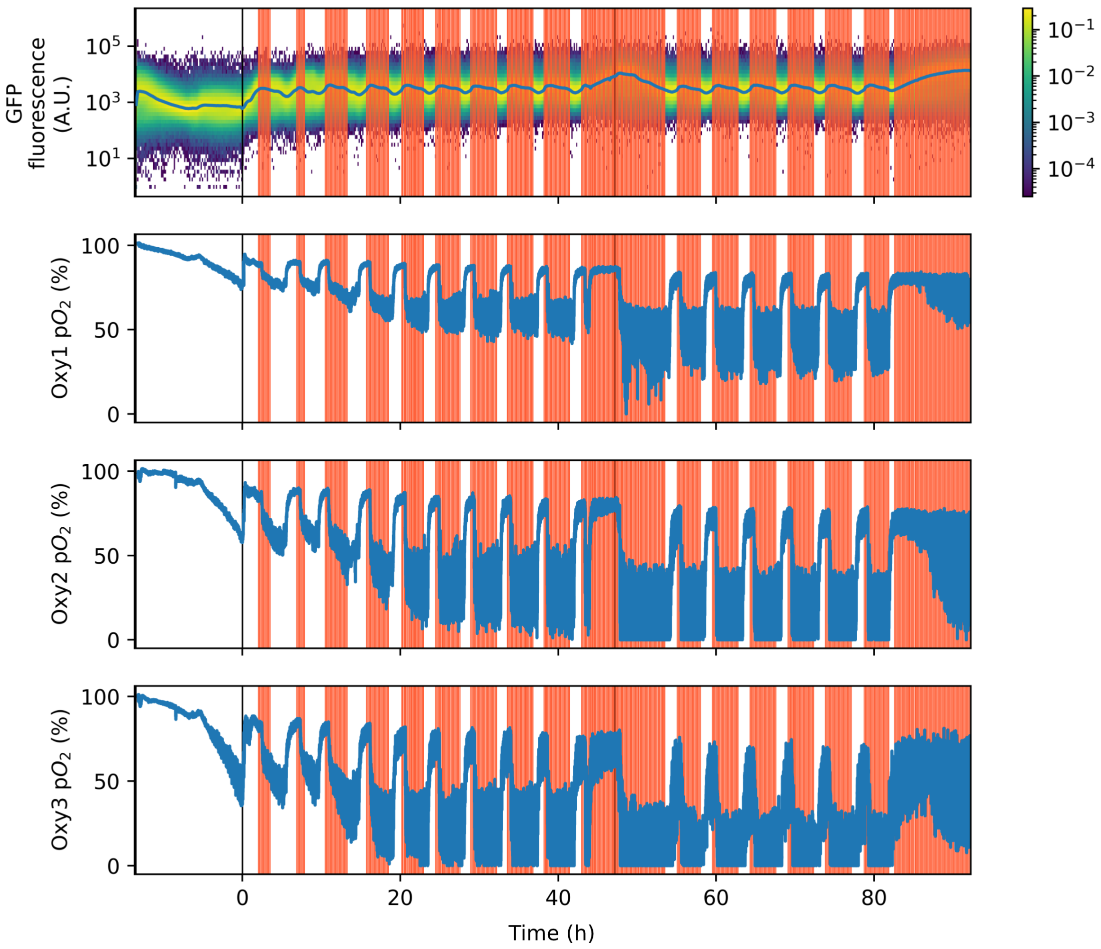


Figure S7: Dissolved oxygen dynamics in the recirculation loop during experiment "2021-06-30_FDE_SD_Seg_Sacch_Tr4-8min." The top panel shows single-cell GFP fluorescence distributions over time with the median indicated in blue. The three lower panels show dissolved oxygen concentrations ($pO_2$, %) measured by three probes positioned along the recirculation loop (Oxy1: 1.25 m, Oxy2: 5 m, both upstream of the feeding point; Oxy3: 8.75 m downstream of the feeding point). The experiment was performed in segregostat mode with a residence time (Tr) of 4 min followed by 8 min. Transitions between conditions (batch phase, segregostat Tr = 4 min, segregostat Tr = 8 min) are indicated by vertical black lines. The final series of pulses in the Tr = 4 min condition does not correspond to true pulses due to depletion of the glucose pulse bottle.

#### 5.1. Extended materials and methods

Measurements of dissolved oxygen in the recirculation loop were performed during scale-down experiments in B-Twin bioreactors (Sartorius) (see 7.1. Bioreactors cultivations). Oxygen was monitored using OXY-4 mini probes (PreSens Precision Sensing GmbH, Regensburg, Germany). Two probes were positioned in the starvation zone (at 1.25 m and 5 m from the beginning of the recirculation loop), and one probe was positioned in the excess zone, downstream of the feeding point (at 8.75 m).

### Supplementary Note 6. CFD simulation of industrial aerated fermentation of *S. cerevisiae*

Examples of cell lifelines from CFD simulation of a $22\text{m}^3$ aerated fermentation of *S. cerevisiae* are displayed in Figure S8.

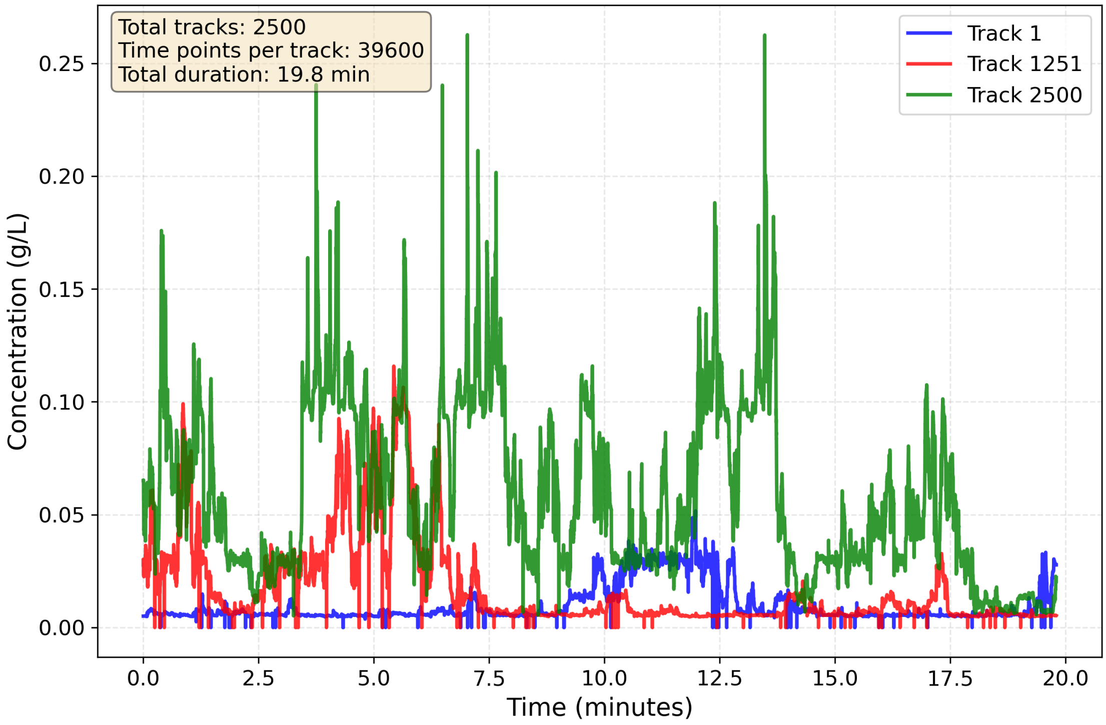


Figure S8: Representative examples of three cell lifelines randomly sampled from the CFD simulation, shown as perceived glucose concentration over time. The 0.03 s timestep ensures proper resolution. Data from Haringa et al. [2].

### Tables

#### 7.1. Bioreactors cultivations

Table 1: Summary of specific operating conditions of the continuous cultures performed in bioreactors (part1).

| ***Culture type and strain*** | ***Operating conditions*** | ***Replicates*** | ***Publication*** |
|---|---|---|---|
| Chemostat (LS)<br>*E. coli* MG1655 $P_{ydcS}$ ::GFPmut2 | Bioreactor : Bionet F1 (Bionet)<br>Cytometer : BD Accuri C6+ (BD Biosciences)<br>D = 0.3 $h^{-1}$ pH = 7 T = 37 °C<br>Aeration = 1 L/min Agitation : 1200 rpm<br>$[glucose]_{FEED}$ = 5 g/L | 2025-01-13_MaD_Chem_Ecoli_ydcS<br>2025-01-29_MaD_Chem_Ecoli_ydcS | This paper<br>Delvenne et al. (2025) |
| Segregostat (LS)<br>*E. coli* MG1655 $P_{ydcS}$ ::GFPmut2 | Bioreactor : Bionet F1 (Bionet)<br>Cytometer : BD Accuri C6+ (BD Biosciences)<br>D = 0.3 $h^{-1}$ pH = 7 T = 37 °C<br>Aeration = 1 L/min<br>Agitation : 1000 & 1200 rpm<br>$[glucose]_{FEED}$ = 5 g/L<br>Regulation threshold = 1000 & 1150 A.F.U.<br>Regulation pulse = 1 g glucose (5.7 ml) | 2024-09-11_FDE_Seg_Ecoli_ydcS<br>2025-01-29_MaD_Seg_Ecoli_ydcS | Delvenne et al. (2025) |
| Chemostat (LS)<br>*S. cerevisiae* CEN.PK 113-7D $P_{glc3}$ ::eGFP | Bioreactor : Bionet F1 (Bionet)<br>Cytometer : BD Accuri C6+ (BD Biosciences)<br>D = 0.1 $h^{-1}$ pH = 5 T = 30°C<br>Aeration = 1 L/min Agitation : 1000 rpm<br>$[glucose]_{FEED}$ = 5 g/L | 2025-02-10_MaD_Chem_Sacch<br>Replicates* | This paper<br>* |
| Segregostat (LS)<br>*S. cerevisiae* CEN.PK 113-7D $P_{glc3}$ ::eGFP | Bioreactor : Bionet F1 (Bionet)<br>Cytometer : BD Accuri C6+ (BD Biosciences)<br>D = 0.1 $h^{-1}$ pH = 5 T = 30°C<br>Aeration = 1 L/min Agitation : 1000 rpm<br>$[glucose]_{FEED}$ = 5 g/L<br>Regulation threshold = 3000 A.F.U.<br>Regulation pulse = 0.4 g glucose (5.7 ml) | 2025-02-10_MaD_Seg_Sacch<br>Replicates* | Delvenne et al. (2025)<br>* |
| Chemostat (SD - Tr = 2min)<br>Segregostat (SD - Tr = 2min)<br>Chemostat (SD - Tr = 8min)<br>Segregostat (SD - Tr = 8min)<br>*E. coli* MG1655 $P_{ydcS}$ ::GFPmut2 | Bioreactor : Bionet F1 (Bionet)<br>Cytometer : BD Accuri C6+ (BD Biosciences)<br>D = 0.3 $h^{-1}$ pH = 7 T = 37 °C<br>Aeration = 1 L/min<br>Agitation : 1200 rpm<br>$[glucose]_{FEED}$ = 5 g/L<br>Tresidence in the loop = 2 min and 8 min<br>Regulation threshold (Seg.) = 1000 A.F.U.<br>Regulation pulse (Seg.) = 1 g glucose (5.7 ml) | 2025-01-06_MaD_SD_ChemSeg_Ecoli_Tr2-Tr8min<br>2025-01-13_MaD_SD_ChemSeg_Ecoli_Tr2-Tr8min | This paper |

* Replicates not provided in this work. They can be found in Henrion et al. (2023) and in Delvenne et al. (2025).

Table 2: Summary of specific operating conditions of the continuous cultures performed in bioreactors (part2).

| ***Culture type and strain*** | ***Operating conditions*** | ***Replicates*** | ***Publication*** |
|---|---|---|---|
| Chemostat (SD - Tr = 8min)<br>Segregostat (SD - Tr = 8min)<br>*S. cerevisiae* CEN.PK 113-7D $P_{glc3}$ ::eGFP | Bioreactor : Bionet F1 (Bionet)<br>Cytometer : BD Accuri C6+ (BD Biosciences)<br>D = 0.1 $h^{-1}$ (+(Chem.) 0.3 h-1) pH = 5 T = 30°C<br>Aeration = 1 L/min Agitation : 1000 rpm<br>$[glucose]_{FEED}$ = 5 g/L<br>Tresidence in the loop = 8 min<br>Regulation threshold (Seg.) = 3000 A.F.U.<br>Regulation pulse (Seg.) = 0.4 g glucose (5.7 ml) | 2025-02-19_MaD_SD_ChemSeg_Sacch_Tr8min<br>2025-03-03_MaD_SD_ChemSeg_Sacch_Tr8min | This paper |
| | Bioreactor : B-Twin (Sartorius)<br>Cytometer : BD Accuri C6 (BD Biosciences)<br>D = 0.1 $h^{-1}$ pH = 5 T = 30°C<br>Aeration = 1 L/min Agitation : 1000 rpm<br>$[glucose]_{FEED}$ = 5 g/L<br>Tresidence in the loop = 8 min<br>Regulation threshold (Seg.) = 5000 A.F.U.<br>Regulation pulse (Seg.) = 0.4 g glucose | 2021-06-30_FDE_SD_Seg_Sacch_Tr4-8min | |
| Segregostat (SD - Tr = 2min)<br>*S. cerevisiae* CEN.PK 113-7D $P_{glc3}$ ::eGFP | Bioreactor : B-Twin (Sartorius)<br>Cytometer : BD Accuri C6 (BD Biosciences)<br>D = 0.1 $h^{-1}$ pH = 5 T = 30°C<br>Aeration = 1 L/min Agitation : 1000 rpm<br>$[glucose]_{FEED}$ = 5 g/L<br>Tresidence in the loop = 2 min<br>Regulation threshold (Seg.) = 5000 A.F.U.<br>Regulation pulse (Seg.) = 0.4 g glucose | 2021-06-20_FDE_SD_Seg_Sacch_Tr2min<br>** | This paper |
| Chemostat (SD - Tr = 4min)<br>*S. cerevisiae* CEN.PK 113-7D $P_{glc3}$ ::eGFP | Bioreactor : B-Twin (Sartorius)<br>Cytometer : BD Accuri C6 (BD Biosciences)<br>D = 0.1 $h^{-1}$ pH = 5 T = 30°C<br>Aeration = 1 L/min Agitation : 1000 rpm<br>$[glucose]_{FEED}$ = 5 g/L<br>Tresidence in the loop = 4 min | 2021-07-15_FDE_SD_Seg_Sacch_Tr4min<br>** | This paper |
| Segregostat (SD - Tr = 4min)<br>*S. cerevisiae* CEN.PK 113-7D $P_{glc3}$ ::eGFP | Bioreactor : B-Twin (Sartorius)<br>Cytometer : BD Accuri C6 (BD Biosciences)<br>D = 0.1 $h^{-1}$ pH = 5 T = 30°C<br>Aeration = 1 L/min Agitation : 1000 rpm<br>$[glucose]_{FEED}$ = 5 g/L<br>Tresidence in the loop = 4 min<br>Regulation threshold (Seg.) = 5000 A.F.U.<br>Regulation pulse (Seg.) = 0.4 g glucose | 2021-06-30_FDE_SD_Seg_Sacch_Tr4-8min<br>2021-07-15_FDE_SD_Seg_Sacch_Tr4min | This paper |
| Chemostat (LS) with pulsed feed<br>*S. cerevisiae* CEN.PK 113-7D $P_{glc3}$ ::eGFP | Bioreactor : B-Twin (Sartorius)<br>Cytometer : BD Accuri C6 (BD Biosciences)<br>D = 0.1 $h^{-1}$ pH = 5 T = 30°C<br>Aeration = 1 L/min Agitation : 1000 rpm<br>$[glucose]_{FEED}$ = 0 g/L<br>Pulse glucose : every 5, 10 or 30 min, from a 20% solution. The total amount of glucose supplied over time was the same as in the chemostat cultivation at D = 0.1 $h^{-1}$. | 2019-07-16_BoZ_Pulsed_Sacch_5min<br>2019-08-13_BoZ_Pulsed_Sacch_5min<br><br>2019-07-09_BoZ_Pulsed_Sacch_10min<br>2020-02-05_BoZ_Pulsed_Sacch_10min<br><br>2019-07-29_BoZ_Pulsed_Sacch_30min<br>2020-02-12_BoZ_Pulsed_Sacch_30min | This paper |

** No replicate available

Table 3: Chronological sequences of operating phases in multi-condition experiments.

| ***Experiment*** | ***Conditions sequences*** |
| --- | --- |
| 2025-01-13_MaD_Chem_Ecoli | Chem. (t = 97:46 , d = 25:49) (following 2025-01-13_MaD_SD_ChemSeg_Ecoli_Tr2-Tr8min) |
| 2025-01-29_MaD_ChemSeg_Ecoli | Batch (t = 0 , d = 5:48) ; Chem. (t = 5:48 , d = 21:06) ; Seg. (t = 45:52 , d = 24:43) |
| 2025-02-10_MaD_ChemSeg_Sacch | Batch (t = 0:0, d = 18:54); Chem. D=0.3 (t = 18:54, d = 27:25);<br>Chem. D=0.1 (t = 46:19, d = 74:38); Seg. D=0.1 (t = 160:09, d = 29:1) |
| 2025-01-06_MaD_SD_ChemSeg_Ecoli_Tr2-Tr8min | Batch (t = -2:35, d = 6:10) ; Chem. Tr=2 (t = 3:35, d = 17:19) ; Seg. Tr= 2 (t = 20:54, d = 27:21) ;<br>Chem. Tr=8 (t = 48:15, d = 19:23 ) ; Seg. Tr= 8 (t = 67:38, d = 23:37) |
| 2025-01-13_MaD_SD_ChemSeg_Ecoli_Tr2-Tr8min | Batch (t = 0, d = 5:27) ; Chem. Tr=2 (t = 5:27, d = 18:20) ; Seg. Tr= 2 (t = 23:47 , d = 25:40 ) ;<br>Chem. Tr=8 (t = 49:27, d = 19:29) ; Seg. Tr= 8 (t = 68:56 , d = 28:50); + LS: 2025-01-13_MaD_Chem_Ecoli |
| 2025-02-19_MaD_SD_ChemSeg_Sacch_Tr8min | Batch (t = 0, d = 18:5); Chem. D=0.1 Tr=8 (t = 18:5, d = 52:27); Chem. D=0.3 Tr=8 (t = 70:32, d = 19:34);<br>Seg. D=0.1 Tr=8 (t = 90:06, d = 76:15) |
| 2025-03-03_MaD_SD_ChemSeg_Sacch_Tr8min | Batch (t = 0, d = 18:32); Chem. D=0.1 Tr=8 (t = 18:32, d = 53:11); Chem. D=0.3 Tr=8 (t = 71:43, d = 22:05);<br>Seg. D=0.1 Tr=8 (t = 93:48, d = 71:53) |
| 2021-06-20_FDE_SD_Seg_Sacch_Tr2min | Batch (t = 0, d = 14:30); Seg. D=0.1 Tr=2 (t = 14:30, d = 85) |
| 2021-07-15_FDE_SD_ChemSeg_Sacch_Tr4min | Batch (t = 0, d = 13:00); Chem. D=0.1 Tr=4 (t = 13:00, d = 46:30); Seg. D=0.1 Tr=4 (t = 59:30, d = 33:30) |
| 2021-06-30_FDE_SD_Seg_Sacch_Tr4-8min | Batch (t = 0, d = 13:45); Seg. D=0.1 Tr=4 (t = 13:45, d = 47:00); Seg. D=0.1 Tr=8 (t = 60:45, d = 35:30) |

t : start time (h), d : duration (h), LS : Lab-Scale, SD : Scale-Down, Chem. : chemostat, Seg. : Segregostat, D : dilution rate ($h^{-1}$), Tr : retention time in the recirculation loop (min)

### Data and Code accessibility

#### 8.1. Data

Raw data are available at https://doi.org/10.5281/zenodo.17453611.

They are separated in 6 folders.

**A_LabScale** :
Contains the raw data from the reactors experiments performed in lab-scale bioreactor.

**B_ScaleDown** :
Contains the raw data from the reactors experiments performed in scale-down bioreactor.

**BZ_Pulses** :
Contains the raw data from the reactors experiments performed in lab-scale bioreactor with feeding by pulses (in Supp. Materials only).

**C_Changestats** :
Contains the raw data from the reactors experiments of accelerostats and decelerostats performed in lab-scale bioreactor (in Supp. Materials only).

**CytometerCalibration** :
Contains the data for calibrations of cytometers (from [4])

**Paper_FEC** :
Contains data published in [4] for the computation of the trade-off between growth and targeted gene expression.

For each cultivation in bioreactor, summary plots are provided together with the raw data. These plots include measured and computed parameters such as fluorescence time-scatter plots from flow cytometry with the corresponding entropy, cell biomass, pH, acid/base addition, temperature, mixing, and dissolved oxygen.

#### 8.2. Code

Relevant code for data analysis and figures assembly is available at https://gitlab.uliege.be/mipi/published-software/2025-scaledownsegregostat.
